# High-resolution genome-wide mapping of chromosome-arm-scale truncations induced by CRISPR-Cas9 editing

**DOI:** 10.1101/2023.04.15.537038

**Authors:** Nathan H. Lazar, Safiye Celik, Lu Chen, Marta Fay, Jonathan C. Irish, James Jensen, Conor A. Tillinghast, John Urbanik, William P. Bone, Genevieve H. L. Roberts, Christopher C. Gibson, Imran S. Haque

## Abstract

CRISPR-Cas9 editing is a scalable technology for mapping of biological pathways, but it has been reported to cause a variety of undesired large-scale structural changes to the genome. We performed an arrayed CRISPR-Cas9 scan of the genome in primary human cells, targeting 17,065 genes for knockout with 101,029 guides. High-dimensional phenomics reveals a “proximity bias” in which CRISPR knockouts bear unexpected phenotypic similarity to knockouts of biologically-unrelated genes on the same chromosome arm, recapitulating both canonical genome structure and structural variants. Transcriptomics connects proximity bias to chromosome-arm truncations. Analysis of published large-scale knockout and knockdown experiments confirms that this effect is general across cell types, labs, Cas9 delivery mechanisms, and assay modalities, and suggests proximity bias is caused by DNA double-strand-breaks with cell cycle control in a mediating role. Finally, we demonstrate a simple correction for large-scale CRISPR screens to mitigate this pervasive bias while preserving biological relationships.

## Introduction

CRISPR-Cas9-based methods are powerful tools for genome editing, with applications ranging from *in vitro* discovery biology, to *ex vivo* editing for cell therapies, and *in vivo* editing for genetic therapeutics [Raguram2022]. Unlike earlier technologies, Cas9 is both programmably scalable—requiring only design of a new guide RNA to target a new gene, rather than a full protein design as with zinc finger nucleases or TALENs—and relatively specific, unlike siRNAs, which are programmable but have broad off-target effects [Jackson2010, Becker2021].

Although CRISPR-Cas9-based editing is much more specific to a chosen target relative to earlier technologies, it is known that guides which are nominally specific to a given genomic position may in fact have off-target activity genomically distant from the intended target [Hsu2013]. Furthermore, evidence has grown to suggest that protocols employing the nuclease activity of Cas9 can lead to undesired on-target changes to the genome *proximal* to the target site, with reports ranging from kilobase-scale deletions [Adikusuma2018, Geng2022], to chromosome truncation or loss [Zuccaro2020, Papathanasiou2021, Tsuchida2023], to more complex rearrangements including duplications and inversions [Kosicki2018, Amendola2022, Geng2022]. Profiling the rate and impact of these effects systematically across the genome is therefore of paramount importance to both discovery and therapeutic development. However, the cost and labor intensity of performing such a screen using existing molecular or sequencing-based methods has practically precluded such a scan.

Cellular morphological profiling, or “phenomics,” is an emerging technology for high-dimensional phenotyping that presents a powerful alternative to transcriptomic or proteomic assays. These microscopy-based methods measure cellular morphology, which is a holistic functional endpoint of cellular state. Like molecular assays, morphological profiling can extract hundreds or thousands of biologically meaningful dimensions of variation from each sample [Bray2016]. However, the use of simple fluorescent stains and imaging, without complicated chemistry or microfluidics, means that morphological profiling is able to generate high-dimensional and intrinsically spatial single-cell data at much lower cost than molecular methods, enabling ultra-large scale perturbative experiments that are uneconomical today with methods like single-cell RNA sequencing.

Here, we apply phenomics to systematically profile CRISPR-Cas9 knockouts of the human genome in primary cells, individually targeting over 17,000 genes with over 100,000 CRISPR guides. Six-stain fluorescent images of plated cells [Bray2016] are encoded into a feature vector using a proprietary deep-learning model, normalized to controls and aggregated across replicate wells, distinct targeting guides and experimental repeats to produce a single “gene vector” mathematically representing the phenotype of each perturbed gene. Cosine similarity between these gene vectors in a high-dimensional representation space acts as a pairwise measure of phenotypic similarity between knockouts recapitulating both known and novel biological relationships including protein complexes and annotated pathways and can be extended to assess similarity among a broad range of cellular perturbations, including genetic, large molecule, and small molecule treatments [Celik2022].

In this work we report a novel observation of “proximity bias”, in which phenotypes of CRISPR knockouts are systematically more similar to the phenotypes of knockouts not only of biologically related genes, but also those of unrelated genomically proximal genes located on the same chromosome arm. Through replication in independent datasets, we demonstrate that this effect is general across laboratories, analysis methods, cell types, and Cas9 delivery mechanisms. Further, we find that the effect is dependent on nuclease activity and not detected in shRNA or CRISPR interference (CRISPRi) experiments. These high-resolution patterns of proximity bias recapitulate chromosomal structure including large-scale structural variants. Molecular investigation of this effect with bulk and single-cell transcriptomic analysis supports large-scale chromosomal truncation in a subpopulation of cells as a driving mechanism. Additionally we reanalyze the DepMap genome-wide CRISPR-Cas9 screens [Behan2019] in cancer cell lines bearing diverse genomic backgrounds to confirm the impact of proximity bias on target discovery and to propose potential mediators for proximity bias. Finally, we show that an arm-based geometric normalization of gene-level features largely corrects for this bias without impacting the recovery of known biological relationships.

## Results

### Phenomics enables high-resolution genome-wide knockout profiling and recapitulates known biological associations

To produce a genome-wide “map” of pairwise phenotypic similarity between gene knockouts, we performed a single-timepoint phenomics screen using CRISPR-Cas9 to knock out 17,065 genes in primary human umbilical vein endothelial cells (HUVEC) with 101,029 guides (typically 6 guides per gene and 24 replicates per guide) leveraging a highly automated robotics-enabled workflow (rxrx3 dataset) [Fay2023].

To further validate the power of phenomics, we additionally computed a complementary similarity map based on the independent cpg0016 dataset from the Joint Undertaking in Morphological Profiling-Cell Painting consortium [Chandrasekaran2023]. There are key differences between rxrx3 and cpg0016. First, the cpg0016 dataset profiles fewer genes (n=7,975) with fewer guides per gene (n=4 guides and 5 replicates per guide), delivered as a pool rather than individually, and screens the U2OS osteosarcoma cell line rather than primary endothelial cells (HUVEC). Second, the Cas9 and guide delivery protocol differs, with the rxrx3 dataset using lipofection delivery of Cas9-guide ribonucleoprotein complexes with one guide per well, whereas the cpg0016 dataset used a Cas9-integrated U2OS cell line (from lentiviral delivery) and lipofection of pools of 4 guide RNAs per well. Third, morphological features of cpg0016 were extracted from CellProfiler [Carpenter2006], whereas a Recursion-proprietary deep learning model was used to extract features directly from the rxrx3 imaging data. For both datasets, gene vectors were aggregated across guides targeting the same gene, intra-batch replicates, and batches to build a genome-wide “map” in which the vector representations of individual gene knockouts could be compared for phenotypic similarity or dissimilarity using cosine similarity (Figure 1a).

**Figure 1:**
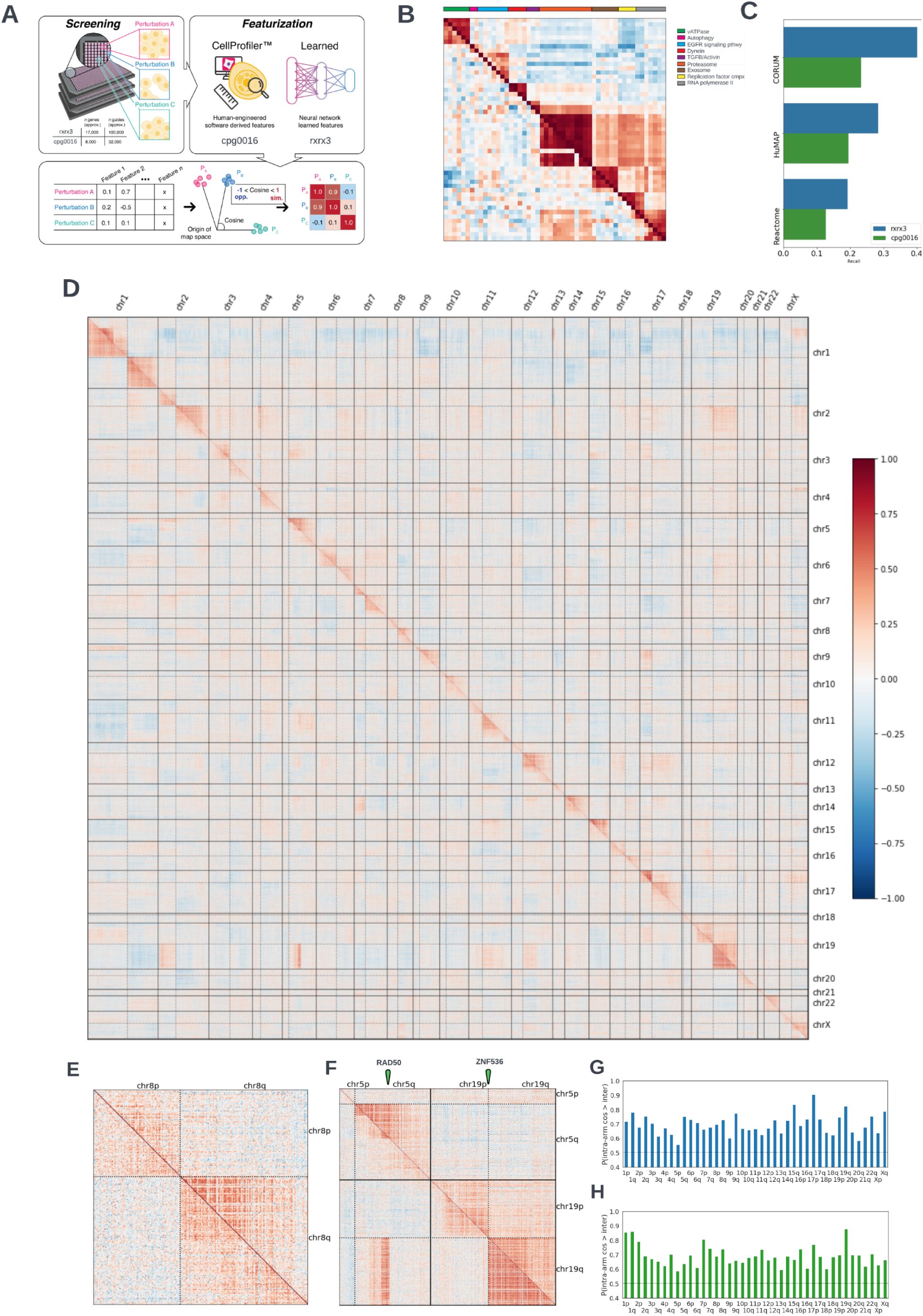
Heatmaps of phenotypic similarity between gene knockouts recapitulate known biology as well as genomic proximity effects. In all heatmaps, each row and column represent a single gene with rxrx3 data shown above the diagonal and cpg0016 data below. Only the 7,477 genes that are present in both datasets are shown. Solid lines represent chromosome boundaries and dashed lines indicate centromeres. A. Phenomics overview. Array-based screening on genetic perturbations produces images of cells which are then quantized either using human-engineered software (CellProfiler) or neural-network learned features. Feature vectors for each pair of perturbations in a high dimensional latent space are related using cosine similarity (ranging from -1 to 1) and visualized in colored heatmaps where each row and column represent a single gene. B. Heatmap of genes with diverse functions demonstrating the ability of phenomics to cluster genes with similar known biological functions. Rows and columns are clustered on the Recursion data. C. Illustrated is the recall of annotated known relationships from each database in the most extreme 10% of similarities (two-sided). Higher values indicate that more relationships in annotation sets are prioritized into the positive and negative tails of the distribution. As a baseline, a random ranking of gene-gene pairs would give a value of 0.1. Not all known biology in any annotation set would be expected to be present in any single cell type. D. Full-genome heatmap where each row and column represent a gene assessed in both rxrx3 (above diagonal) and cpg0016 (below diagonal) studies. Ordering genes by chromosomal position reveals the proximity bias signal along the diagonal present in both datasets. Chromosome boundaries are clearly visible (solid lines) as are centromere locations (dotted lines). E. A zoom in on chromosome 8 showing that proximity bias, while visible at large scales, does not impact all genes equally. F. Juxtaposition of chromosomes 5 and 19, where the pattern of proximity bias signal reflect a chromosomal rearrangement known to be present in U2OS cells (cpg0016 data) but not HUVEC (rxrx3 data). G. Barplot of proximity bias metrics (Brunner-Munzel probabilities) for each chromosome arm for the rxrx3 dataset. Values above 0.5 indicate elevated intra-arm similarity, and all chromosome arms are highly significant with Bonferroni correction (p < 0.001). H. Barplot of proximity bias metrics (Brunner-Munzel probabilities) for each chromosome arm for the cpg0016 dataset. Values above 0.5 indicate elevated intra-arm similarity, and all chromosome arms are highly significant with Bonferroni correction (p < 0.001).

We evaluated the ability of rxrx3 and cpg0016-based maps to recapitulate known biology in both a targeted and a broad sense. Targeted examination of genes in well-studied pathways showed that phenomics-derived gene-gene similarities recapitulate both biology that is highly conserved across cell types (e.g., microtubule, proteasome, and autophagy genes), as well as therapeutically-relevant pathways including JAK/STAT, TGF-beta, and insulin receptor) (Figure 1b, Figure S1). Despite the methodological differences between datasets, large-scale benchmarking [Celik2022] of both datasets shows substantial recall of known annotations drawn from public datasets including Reactome [Gillespie2022], HuMAP [Drew2021], and CORUM [Giurgiu2019] (Figure 1c).

### CRISPR-Cas9 knockouts are systematically more similar to those on the same chromosome arm, revealing canonical and variant genome structure

Upon the generation of the rxrx3 full-genome knock-out data, we noticed a curious bias: the distribution of cosine similarities for relationships between genes on the same chromosome was shifted relative to the distribution of similarities for gene pairs on different chromosomes (Figure S2a). To explore this phenomenon further, we visualized the full genome-wide dataset ordered by genomic coordinate, revealing a striking structure in which knockouts of genomically proximal genes were systematically more phenotypically similar to one another than to knockouts of genes on other chromosomes (Figure 1d,e above diagonal). Closer examination reveals that these systematic blocks of “proximity bias”, in which phenotypic similarity is correlated with genomic proximity of the genes rather than biological relevance, are terminated by centromeres, indicating chromosome-arm-scale effects (Figure 1d,e above diagonal).

To test whether proximity bias was an artifact of our laboratory’s protocol, our Cas9 and guide delivery system was specific to HUVEC or a consequence of our computational image analysis and featurization scheme, we performed the same visualization in cpg0016 (Figure 1d,e below diagonal). The proximity bias observed in rxrx3 was also evident in cpg0016 both visually in the genome-wide heatmap and in intra- vs inter-chromosomal similarity distributions (Figure S2a).

As proximity blocks appeared to correlate with chromosomal structure, and the rxrx3 dataset screens were performed in karyotypically-normal human primary cells, we wondered whether proximity blocks would also reflect non-reference structure in genomically abnormal cells. The U2OS line used in cpg0016 is known to be karyotypically abnormal and heterogeneous, with different clones exhibiting distinct genotypes [Raftopoulou2020]; a number of known fusion events have been cataloged in U2OS at DepMap [Tsherniak2017], including a fusion between *RAD50* on chromosome 5q and *ZNF536* on chromosome 19q. Examination of the cpg0016 map shows a clear block of inter-chromosomal proximity bias between 5q and 19q that closely recapitulates the boundaries of the annotated fusion (Figure 1f).

Finally we quantified the proximity bias effect by estimating the probability, for each arm, of a within-chromosome arm relationship displaying a higher cosine similarity than a between-chromosome arm relationship using a non-parametric Brunner-Munzel test [Brunner2000]. This metric is both comparable across maps that may have different numbers of tested genes, and flexible enough to be used to quantify the bias encoded in an entire map (comparing the full set of similarities), within each chromosome arm, or at the gene level (by restricting to relationships involving that arm or gene). At the full-map level, the probability of a within-arm relationship being ranked above a between-arm relationship was 0.71 for the rxrx3 dataset and 0.72 for the cpg0016 data (p-values < 0.001). At the chromosome-arm level, both datasets show a significant effect for all chromosome arms (Figure 1g,h).

### Proximity bias arises from widespread chromosome arm truncation

A number of potential unintended target-proximal consequences of Cas9 editing have been discussed in the literature, ranging from kilobase-scale deletions, translocations, chromothripsis, complete or partial chromosome loss, and complex rearrangements involving deletion, duplication, and inversion [Kosicki2018, Adikusuma2018, Alanis-Lobato2021, Cullot2019, Leibowitz2021, Amendola2022, Papathanasiou2021, Barkal2016, Zuccaro2020, Geng2022, Przewrocka2020, Weisheit2020, Tsuchida2023]. Although the rxrx3 and cpg0016 genome-wide maps present a compelling case for widespread proximity bias, they do not directly nominate a mechanism.

Close examination of the genome-wide maps in Figure 1 demonstrates a recurrent pattern in which knockouts of genes closer to a centromere display a stronger proximity bias signal. To rigorously assess this observation, we plotted the gene-level Brunner-Munzel statistic versus the relative chromosome arm position for both datasets and calculated the Spearman correlation (Figure 2a and Figure S2b). Most of the chromosome arms showed a significant correlation in both datasets, and these values agree well with the fading intensity of blocks in full-genome heatmap visualization (Figure 2b and Figure S2c). This correlation, combined with the recovery of known biology already established in these maps, suggested a model illustrated in Figure 2c. In this model, specific knockouts of particular genes can produce unique phenotypes (allowing recall of on-pathway biology in the unconfounded case), but Cas9 editing occasionally results in chromosomal truncations deleting large chromosomal segments from the intended cut site towards the telomere. These truncations delete *multiple* genes that would individually exhibit phenotypes, thus producing a mixed phenotype modeled as the sum of the individual phenotypes. Such a model induces a hypothesis testable by molecular methods.

**Figure 2:**
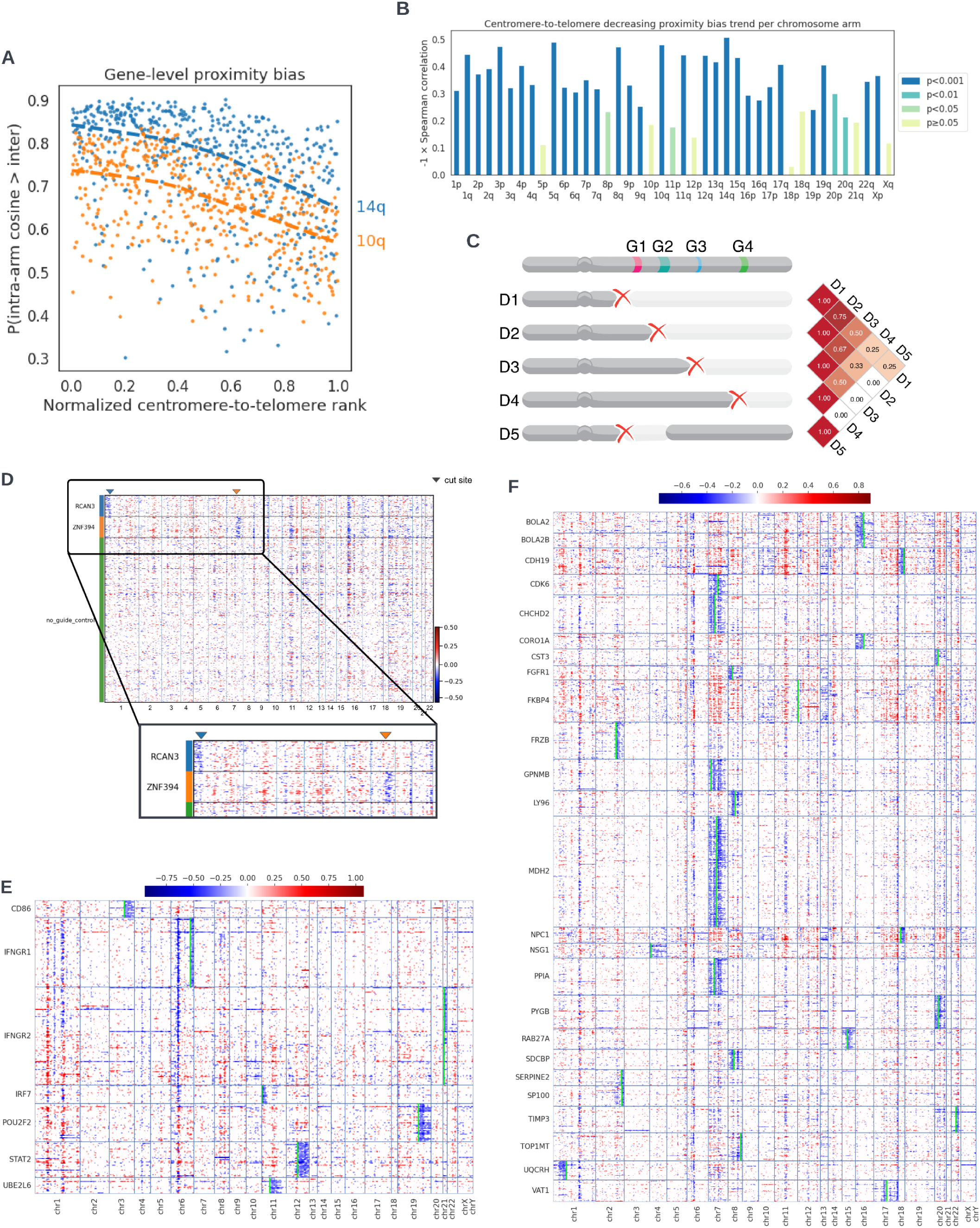
Genome-wide phenomic measurements and transcriptomic data support model of chromosomal truncation underlying proximity bias. A. Scatter plot of gene-level proximity bias metric vs position with the chromosome arm for two chromosome arms. A negative correlation supports the visual pattern seen in figure 1C of weakening proximity bias signal toward telomeres. B. Spearman correlations in plots similar to A across all chromosome arms. The height of the bar for each arm agrees well with the degree of fading in diagonal blocks in Figure 1D. P-values show significance of the correlation. C. Schematic representation of how the pattern of weakening signal toward telomeres observed in chromosome-arm heatmaps may arise by deletions sharing loss of function of multiple key genes. D1,…,D4 represent varying-length truncations and D5 an intra-arm deletion. D1 and D2 both lose three key genes and hence are highly similar, while D1 and D4 only share one and so may look less similar. The intra-arm deletion D5 only shares one key gene loss with D1 and zero losses with D2-D4 and therefore is expected to show less proximity bias. D. Heatmap of CNV estimates in bulk RNA-seq samples showing loss of chromosome regions telomeric to the cut site of guides targeting introns in the RCAN3 and ZNF394 loci. Each row in the heatmap represents a treatment well. Each column represents a gene block, ordered by the chromosome number. *Heatmaps of CNV estimates in two scRNAseq datasets for genes exhibiting deletions near the cut site when targeted by CRISPR-cas9. Each row in each heatmap represents a cell exhibiting deletion near the target gene site as in the row label. The rows are ordered alphabetically based on target gene name. Each column represents a gene block, ordered by the chromosome number. Lime bars represent the target gene sites*. E. CNV estimate heatmap for data from Papalexi et al., 2021. F. CNV estimate heatmap for data from Frangieh et al., 2021.

Similar truncations and deletions have been previously observed in the literature as rare consequences of Cas9 editing (e.g., a single-cell analysis of T cells edited at the *TRAC* gene suggested about 3% of cells suffered partial or total chromosome loss [Tsuchida2023]). We searched an internal Recursion database of bulk RNA-seq data from HUVEC, comprising 25,564 samples with an average of 1.3M unique reads per well. To enhance the sensitivity of our analysis, we focused on a high-replicate subset, in which 45 genes were each targeted with a single intronic guide across 63 replicates per gene (total N=2,835 samples), compared against a no-guide reference pool of 3,320 samples. Comparing copy number variation (CNV) events from Cas9-edited wells relative to Cas9-free controls, we observed multiple loci enriched for deletions between the target cut site and the telomere, including the genes *ZNF394* (located on chromosome 7q) and *RCAN3* (on chromosome 1p) (Figure 2d, Figure S2d, Table S1). Although the number of loci found in this search was limited, these events are expected to be rare and therefore difficult to detect in bulk assays, particularly by RNA rather than DNA sequencing. Identification of edits with deletions therefore corroborates the model suggested from genome-wide phenomics data that Cas9-induced chromosomal truncations are not locus-specific phenomena, but rather are pervasive across the genome.

**Table 1:** Single-cell sequencing reveals widespread on-target proximal deletions from CRISPR-Cas9. Number and percent of target genes showing deletions specific to the target region in two CRISPR-cas9 datasets uniformly reprocessed in scPerturb.

| Perturbation type | Dataset | Cell type | Total # tested targets | Tested loss direction | % targets w/ specific loss | # targets w/ specific loss | # targets w/ loss towards telomere | # targets w/ loss towards centromere |
| --- | --- | --- | --- | --- | --- | --- | --- | --- |
| CRISPR-Cas9 | Papalexi | THP1 (monocytic leukemia) | 24 | 3' | 33.3 | 8 | 6 | 2 |
|  |  |  |  | 5' | 8.3 | 2 | 1 | 1 |
|  | Frangieh | Melanocytes (melanoma) | 237 | 3' | 10.1 | 24 | 18 | 6 |
|  |  |  |  | 5' | 15.2 | 36 | 21 | 15 |

To further assess whether these molecular results were unique to our CRISPR or transcriptomic protocols, we re-analyzed two datasets uniformly reprocessed as part of the scPerturb resource [Peidli2022] that used single-cell RNA sequencing to examine the effects of CRISPR-Cas9 gene knockout in the THP1 leukemia line and in melanoma-derived melanocytes [Papalexi2021, Frangieh2021]. To address dropout in single-cell sequencing data, we assessed the fraction of targeted genes in each dataset in which substantial, specific loss was observed of the 150 genes closest to the target in either the 3’ or 5’ direction. Across these datasets, 8-33% of targeted genes exhibited copy number loss proximal to the target, further confirming the cell type independence of proximity bias and its root in recurrent deletions or truncations (Table 1). Echoing the results from the bulk RNA sequencing analysis, even for targets with called loss, only a fraction of cells exhibited losses (mean 4.5%, max 13%) (Table S2). As shown in Figure 2e-f, the deletions in called genes from scRNAseq support the proposed model of chromosomal truncation underlying proximity bias.

**Table 2:** Low deletion rates are observed in single-cell RNA sequencing analysis of CRISPRi perturbation datasets. Number and percent of target genes showing deletions specific to the target region in three CRISPRi datasets uniformly reprocessed in scPerturb.

| Perturbation type | Dataset | Cell type | Total # tested targets | Tested loss direction | % targets w/ specific loss | # targets w/ specific loss | # targets w/ loss towards telomere | # targets w/ loss towards centromere |
| --- | --- | --- | --- | --- | --- | --- | --- | --- |
| CRISPRi | Replogle | RPE1 | 2067 | 3' | 1.7 | 36 | 25 | 11 |
|  |  |  |  | 5' | 2.1 | 44 | 13 | 31 |
|  | Tian | iPSC-derived neurons | 177 | 3' | 1.1 | 2 | 1 | 1 |
|  |  |  |  | 5' | 3.4 | 6 | 2 | 4 |
|  | Adamson | K562 (leukemia) | 78 | 3' | 1.3 | 1 | 1 | 0 |
|  |  |  |  | 5' | 0.0 | 0 | 0 | 0 |

### Proximity bias confounds therapeutic target identification

A key application of genome-wide knockout screening is in mapping of biological pathways, particularly for therapeutic target discovery. Consequently, we investigated the potential impact of proximity bias on target discovery in a widely-used publicly available resource.

Project Achilles has performed genome-wide CRISPR screens of cell survival in 625 cancer cell lines in an effort to identify potentially druggable essential genes for a range of tumor types, contributing to the Cancer Dependency Map (DepMap) [Tsherniak2017]. We surmised that if DepMap CRISPR screens were also affected by proximity bias, then it would manifest as patterns of essentiality (across cell types) that cluster unexpectedly by genomic proximity.

To carry out this analysis, we built genome-wide maps from DepMap CRISPR 19Q3 data; in these maps, each gene was characterized by a vector representing its essentiality in each of the 625 tested cell lines, rather than as a vector of morphological features, as in rxrx3 and cpg0016 (Figure 3a). Visual examination and quantification of the DepMap CRISPR map confirms the presence of arm-scale proximity bias (Figure 3b,c). Analysis of a newer version of this data (22Q4), which used more sophisticated copy number and batch correction methods and expanded the number of cell lines to 1,078, also exhibited proximity bias (Figure S3a).

**Figure 3:**
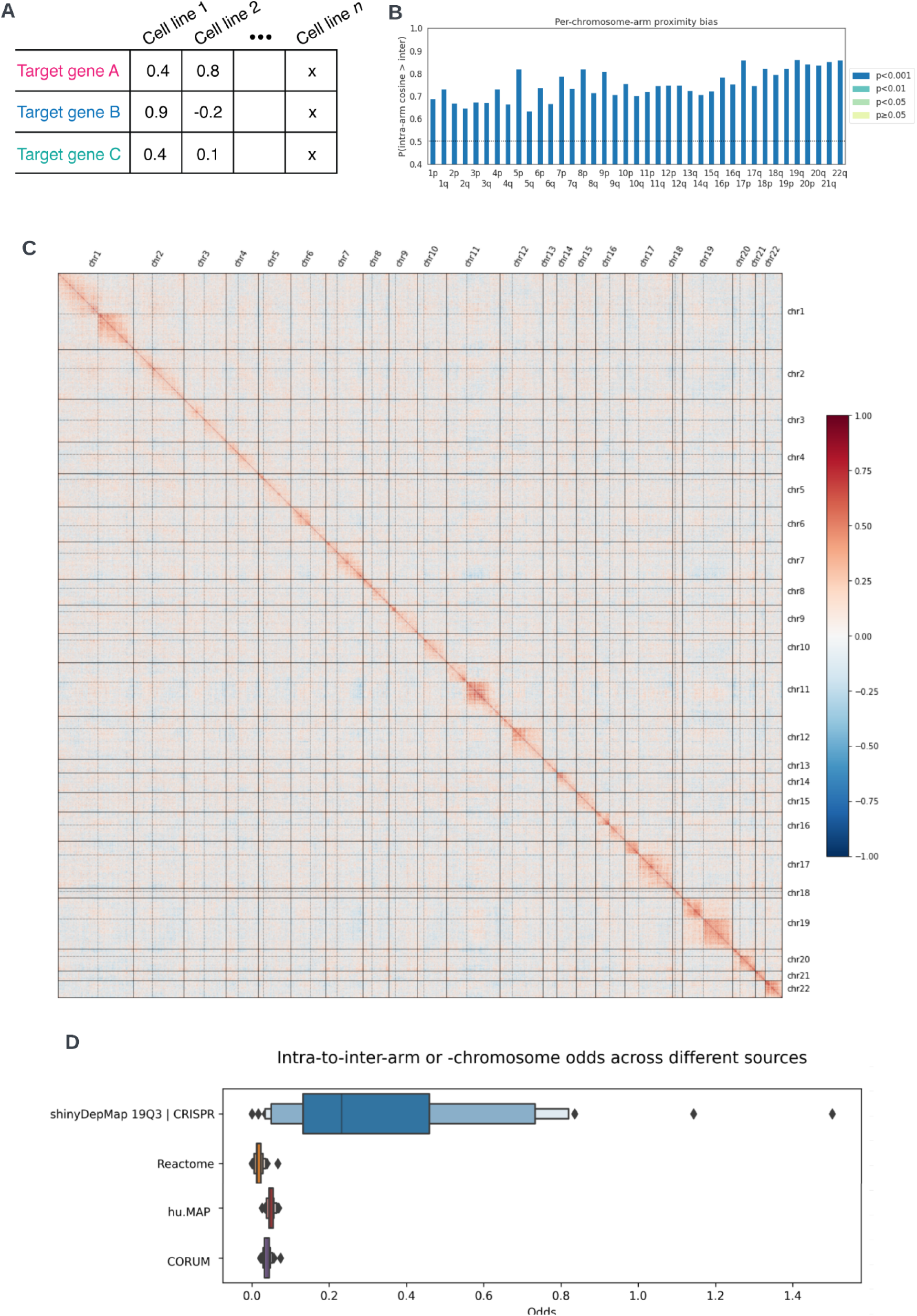
CRISPR-Cas9 screens of cancer gene essentiality are significantly confounded by proximity bias. A. Schematic of the table of feature data used in the DepMap analysis. Each row corresponds to a target gene and each column a cell line. Values in the table are dependency measures of the survival sensitivity for the cell line when the target gene is knocked out with CRISPR-Cas9. We assess similarity between targets by calculating the cosine similarity between rows. B. Barplot of proximity bias metrics (Brunner-Munzel probabilities) for each chromosome arm for the DepMap 19Q3 dataset. Values above 0.5 indicate elevated intra-arm similarity, and all chromosome arms are highly significant with Bonferroni correction. C. Pairwise cosine similarity between DepMap (19Q3) targets ordered by chromosome position across the human genome and quantile normalized to a normal distribution with mean 0 and standard deviation 0.2. Solid lines represent chromosome boundaries and dashed lines indicate centromeres. D. Boxen plots showing distributions of the ratio of within-chromosome arm relationships to between-arm relationships for each chromosome arm across different gene annotation sets. The DepMap data shows a much higher ratio of within-arm to between arm annotations suggesting a systematic bias to the predicted associations.

We next sought to assess the impact of this bias on target discovery. As early stage discovery programs are typically not disclosed by pharmaceutical companies and negative results tend to be underreported in the literature, we assessed whether published DepMap analyses might be confounded by proximity bias. In particular, we examined the results from shinyDepMap [Shimada2021], which sought to cluster genes with similar dependencies to identify druggable targets and pathways using the 19Q3 data.

Examination of 16,941 inferred gene-gene relationships from shinyDepMap reveals that a large number of putative relationships inferred from DepMap CRISPR data are within chromosomal arms. To assess the biological plausibility of this result, we compared the odds of identifying intra- versus inter-arm connectivity in the shinyDepMap clusters against the odds of finding such connectivity in other databases of known biological relationships derived from pathway or protein complex data. shinyDepMap contains far more intra-arm annotated relationships than expected versus this biological prior (Fisher exact test odds ratio = 0.068, p < 0.0001), suggesting that results of downstream DepMap CRISPR analyses are significantly confounded by proximity bias (Figure 3d, Figure S3b).

### Proximity bias is dependent on Cas9 nuclease activity

Given the proposed model of large truncations, we sought to confirm whether proximity bias is dependent on nuclease activity of Cas9. Two non-nuclease dependent technologies for large-scale genetic screens are CRISPR interference (CRISPRi), which uses a mutant Cas9 (optionally fused to a transcriptional repressor) to suppress transcription of target genes [Larson2013], and short-hairpin RNA (shRNA) screens applying RNA interference to knock down target transcripts [Paddison2002]. Comparative analysis of shRNA or CRISPRi knockdown versus CRISPR-Cas9 knockout data can therefore facilitate identifying whether or not nuclease activity is critical to proximity bias. Moreover, comparison of CRISPRi and CRISPR-Cas9 data can address the question of whether other Cas9-specific but non-nuclease-dependent activities also result in proximity bias.

We first extended our analysis of single-cell RNA sequencing CRISPR-Cas9 datasets from scPerturb to three CRISPRi datasets [Replogle2022, Tian2021, Adamson2016]. In contrast to CRISPR-Cas9-perturbed cells, in which 8-33% of assessed genes exhibited large target-site-proximal deletions spanning 150 genes from the perturbed gene start site (Table 1), only up to 3% of target genes were observed to have such losses across three CRISPRi datasets (Table 2, Table S2), suggesting that the nuclease activity of Cas9 is required to generate the elevated proximal deletion rate hypothesized to underlie proximity bias.

We further tested whether the proximity bias observed in DepMap gene essentiality data was a function of Cas9 nuclease activity using DepMap essentiality data derived from shRNA screens. A map built from DepMap shRNA screening data did not exhibit significant proximity bias, suggesting that proximity bias arises as a specific consequence of CRISPR-Cas9 editing, and not as a biological effect arising from knockdown of the targeted genes in cancer cell lines (Figure S5a,b).

### Genetic stratification of DepMap suggests a connection between cell cycle and the generation of proximity bias

We next attempted to elucidate potential biological mechanisms behind and mediators of proximity bias and its associated chromosomal truncations. The DepMap dataset profiled a wide range of cell lines with diverse genetic backgrounds. We therefore hypothesized that it would be possible to stratify cell lines in DepMap by whether they had normal, amplified, or decreased function in particular genes, and to test such stratified splits for changes in proximity bias to identify potential biological mechanisms behind the genesis or suppression of proximity bias.

*TP53* expression has been suggested as a marker for reduced chromosomal loss in CRISPR-Cas9 editing [Cullot2019, Tsuchida2023]; thus, loss of *TP53* would be expected to increase proximity bias by increasing chromosomal loss. As *TP53* is highly mutated in cancer lines, as a first test we stratified DepMap cell lines by *TP53* loss-of-function status, and found significantly increased proximity bias in a CRISPR map built from putatively *TP53*-null cell lines compared with one built using only cell lines with putatively functional *TP53* (Figure 4a).

**Figure 4:**
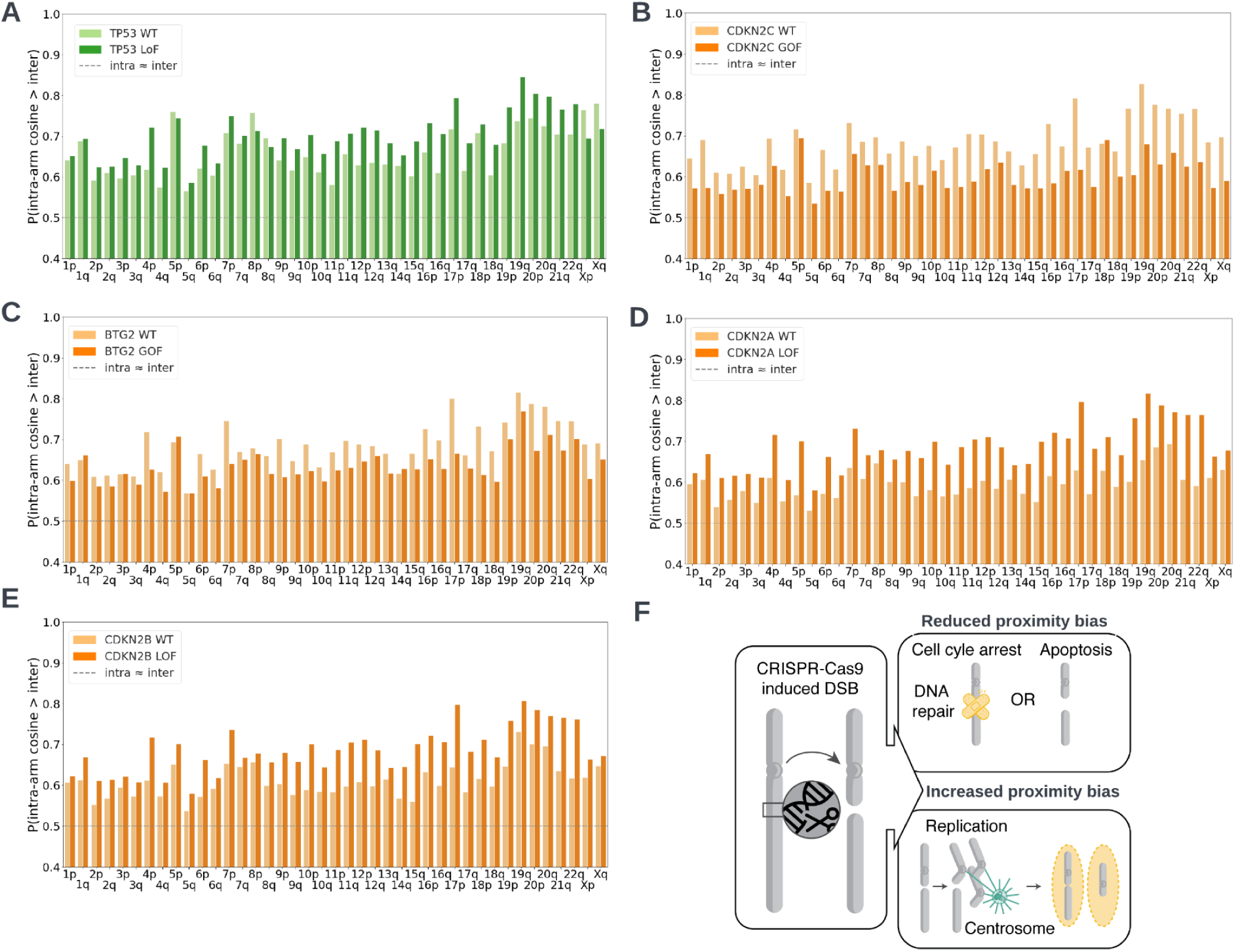
Role of TP53, cell-cycle and replication associated genes in proximity bias. A. Barplot of chromosome arm-level proximity bias quantification by Brunner-Munzel test statistic in *TP53* wild type and *TP53* loss of function cell lines from the DepMap 22Q4 data. Most chromosome arms show increased proximity bias in the loss of function cell lines. *Barplots of chromosome arm-level proximity bias quantification by Brunner-Munzel test statistic in TP53 loss-of-function cell lines from DepMap 22Q4 data. Detailed test statistics in Table S3*. B. Cell lines with amplified *CDKN2C* show reduced proximity bias in most chromosome arms. C. Cell lines with amplified *BTG2* show reduced proximity bias in most chromosome arms. D. Cell lines with loss-of-function of *CDKN2A* show increased proximity bias. E. Cell lines with loss-of-function of *CDKN2B* show increased proximity bias. F. Schematic model for how CRISPR-Cas9 induced double-strand breaks can result in large telomeric deletions. In the presence of a functional p53 protein or other protective factors, cells with double-strand breaks are arrested until the genetic lesion can be repaired or the cell undergoes apoptosis. In the absence (or reduced abundance) of functional p53 protein, a double-strand break may not be repaired before mitosis. As chromatids are segregated through attachment of the centrosome to the kinetochore at the centromere, large segments of DNA stretching from the cut site toward the telomere may fail to segregate and therefore can be degraded or lost in subsequent cell generations.

Having confirmed the connections between *TP53* loss and proximity bias, we attempted to search for additional mechanisms that might mediate proximity bias. We searched DepMap across eight splits: for genes in which putative loss-of-function or amplification either amplified or suppressed proximity bias, in either a *TP53* null or functional background (Table S3). Although DepMap cell lines collectively bear loss-of-function or amplification variants across many genes, individual genes may only be mutated in a small number of lines, and mutation statuses across genes are often highly correlated with each other and with tumor-of-origin. This statistical confounding makes confident estimates of effect for individual genes challenging, as certain genes may be identified spuriously as a consequence of co-occurrence with true positive genes in the same cell lines, either locally in the genome (as a consequence of copy number change) or far apart. Consequently, rather than analyze individual genes beyond *TP53*, we sought signal in enriched biological pathways in each of these contexts (Table S3).

Interestingly, “Negative Regulation of G1/S Transition of mitotic cell cycle” surfaced as an enriched biological pathway in a *TP53* loss-of-function background, based on decreased proximity bias observed in cell lines with increased copy number of *CDKN2C*, *MDM4*, or *BTG2* (Figure 4b,c, Figure S4). *CDKN2C* and *BTG2* inhibit cell cycle progression [Sherr1999, Kim2020] and this data is consistent with the hypothesis that loss of *TP53* and cell cycle progression promotes increased proximity bias. Also, consistent with this hypothesis, loss of the cell cycle inhibitors *CDKN2A* and *CDKN2B* shows increased proximity bias (Figure 4d,e) [Sherr1999]. *MDM4* and *MDM2* act as a heterodimer to regulate p53 and in the absence of *TP53*, *MDM4* and *MDM2* promote cell cycle progression through p73 [Klein2021]. Data we identified here suggests *MDM4* has an opposite effect, but it is unclear whether this represents confounding or novel biology. Beyond cell cycle, we observed enrichment in proliferation pathways and pathways/processes enriched in particular cancer subtypes, which are likely confounded factors based on the cancer background; as well as enrichment in histone modifiers, which are difficult to interpret due to the likely trans-acting consequences of these genes.

These data suggest a possible mechanistic model for the generation of chromosomal truncations by CRISPR-Cas9 leading to proximity bias (Figure 4f). In cells with functional *TP53,* double-strand breaks created by Cas9 are often recognized by p53, leading to cell cycle pause and DNA repair or apoptosis. Without functional p53, mitosis may proceed prior to the repair of the double strand break, leading to daughter cells bearing truncated chromosomes. However, in this scenario, upregulation of other genes (such as *CDKN2C* or *BTG2*) that pause cell cycle may reduce the formation of truncation-bearing cells either through simple mitotic arrest, or by allowing time for repair to proceed. In particular, this model could explain why deletions are often observed from the cut site to the telomere, rather than deletions of the remainder of the chromosome through the centromere, and similarly why proximity bias effects terminate at centromere boundaries: after double-strand break induction, only the chromosomal fragment bearing a centromere can propagate through mitosis.

### Geometric correction reduces proximity bias without impacting recapitulation of known biological relationships

Given that the proximity bias effect appears to be largely localized within chromosome arms, we hypothesized that applying a chromosome-arm correction to rxrx3 and cpg0016 phenomic maps might mitigate the unwanted signal. To that end, we adjusted the vector representation for each gene by subtracting an estimated representation of the chromosome-arm in which the gene is located (see methods). This simple adjustment significantly reduces the proximity bias effect, both globally and per chromosome arm (Figure 5a,b,c,d) while maintaining or improving genome-wide benchmarking metrics in both datasets (Figure 5e). Interestingly, the recall of annotated within-arm relationships decreases with the chromosome-arm correction (Figure 5e center), but this is outweighed by improved recall on a larger number of between-arm annotated relationships, suggesting that the proximity bias effect can confound such benchmarking efforts if it is not taken into account.

**Figure 5:**
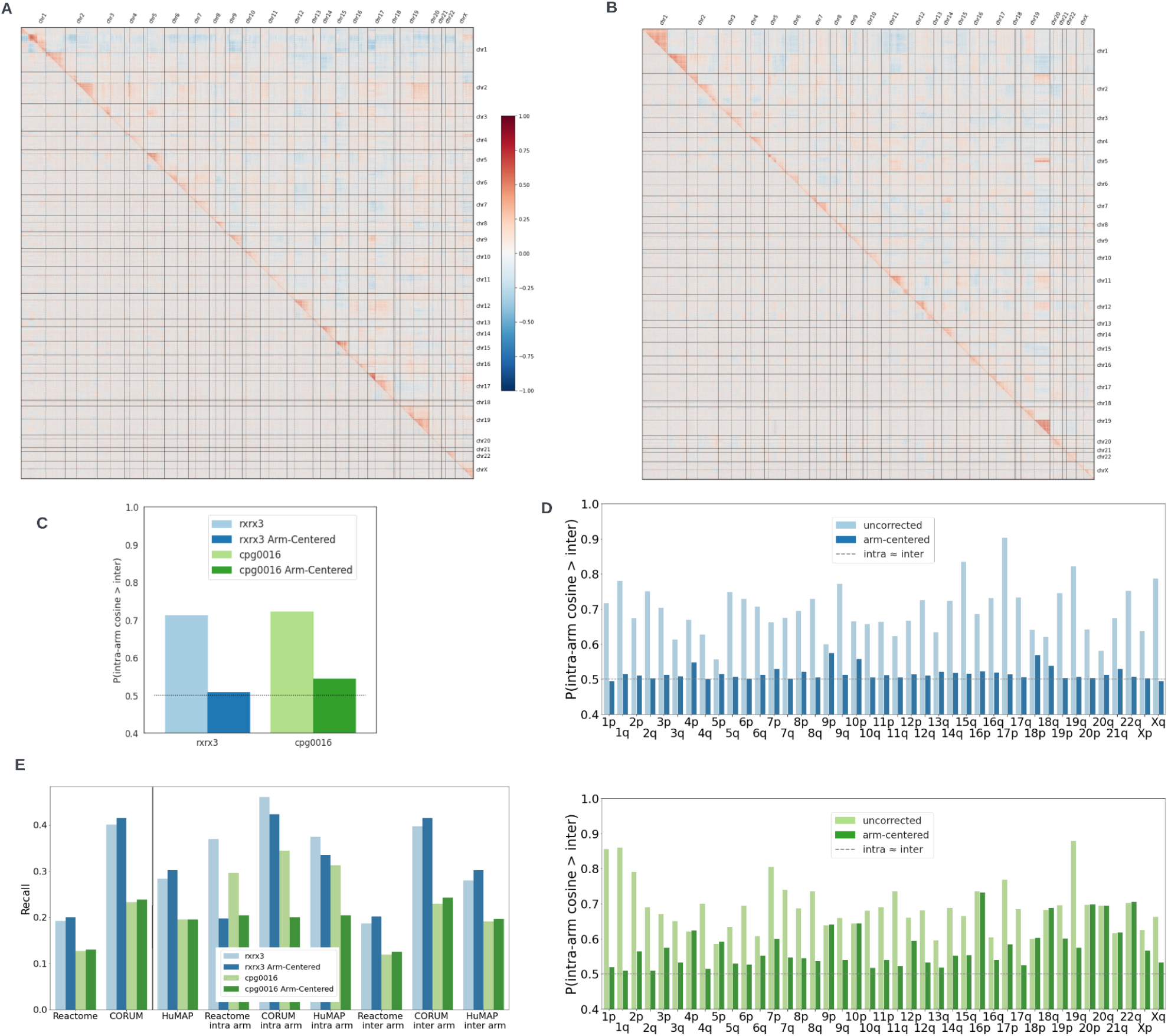
Geometric correction reduces proximity bias effect. *Split heatmaps of pairwise cosine similarities before (above diagonal) and after (below diagonal) chromosome-arm correction*. A. Split heatmap for rxrx3 B. Split heatmap for cpg0016 C. Barplot of genome-wide proximity bias quantification by Brunner-Munzel test statistic before and after chromosome arm correction for rxrx3 and cpg0016. The correction greatly reduces the probability of within-arm relationships having a stronger cosine similarity than between-arm relationships. D. Barplot of chromosome arm-level proximity bias quantification by Brunner-Munzel test statistic before and after chromosome arm correction for rxrx3 (top) and cpg0016 (bottom). The correction reduces the probability of within-arm relationships having a stronger cosine similarity than between-arm relationships for most chromosome arms. E. Annotated relationship recall benchmarking metrics with and without chromosome-arm correction, both at the global-scale (left of vertical line) and stratified by within-arm or between-arm relationships.

## Discussion

Since its discovery, the CRISPR-Cas9 editing system has become an invaluable research tool thanks to its general applicability and programmability, but deeper investigation of this system particularly in the context of gene therapy has suggested potential caveats arising from undesired on-target effects. Because of both the context of their discovery – therapeutic applications examining individual genes – and the expense and labor-intensity of profiling these effects by sequencing, FISH, or other molecular methods, how general these effects are across genomic positions or cellular contexts has not previously been assessed. In this work, we have made use of the scale afforded by cellular phenomics to systematically profile CRISPR-induced gene knockouts for virtually all human protein-coding genes – over 17,000 genes and 100,000 guide sequences – in a primary human cell type, and replicated our findings across cell types, assay contexts, and in molecular follow up.

Three key contributions to the field arise from this analysis. First, undesired on-target proximal effects are not oddities of individual loci in the genome or peculiar to genes of interest in gene therapy, but rather are nearly universal across the genome and likely arise from chromosomal truncations. Second, as a consequence of these effects, possibly all CRISPR-based phenotypic screens, whether based on imaging, sequencing, or functional (e.g. viability) readouts, are confounded by “proximity bias” leading to false discoveries, but it is possible to largely mitigate this by the use of appropriate controls. Finally, these effects are not unique to cancer cell lines, embryos, or embryonic stem cells, but generalize across cell types including normal somatic cells, and therefore may pose safety challenges for *in vivo* gene therapy.

Our main finding is that CRISPR-Cas9-induced truncations are likely to be pervasive across the genome. While a broad range of unintended on-target consequences of Cas9 editing have been discussed in the literature ranging from kilobase-scale deletions, translocations, chromothripsis, complete or partial chromosome loss, and complex rearrangements involving deletion, duplication, and inversion [Kosicki2018, Adikusuma2018, Alanis-Lobato2021, Cullot2019, Leibowitz2021, Amendola2022, Papathanasiou2021, Barkal2016, Zuccaro2020, Geng2022, Przewrocka2020, Weisheit2020, Tsuchida2023], these previous studies have only examined these effects on small numbers of loci (up to hundreds) due to experimental limitations. Here, we leverage the scale and high dimensional readout from phenomics to systematically profile the effects of Cas9 editing across the genome and find that these undesired on-target effects are neither locus-specific nor cell type-specific, but universal across the human genome. The generality of our findings suggests caution in the application of CRISPR-Cas9 knockout editing regardless of genomic location.

Refining this finding of genome-wide generality, our exploration of transcriptional data suggests proximity bias is driven by truncations extending toward telomeres and typically impact a fraction of cells. Prior work has asserted whole-chromosome loss as a potential consequence of CRISPR-Cas9 editing in T cells [Tsuchida2023]. The targeted locus in this work was *TRAC*, located at 14q11.2, near the centromere of the acrocentric chromosome 14. Reanalyzing ECCITE-seq data from THP-1 cells [Papalexi2021], we find evidence for similar occasional loss of the short *p* arm of acrocentric chromosome 21 arising from editing of *IFNGR2* located at 21q22.11. However, in our broader analysis, truncation effects in transcriptomics are almost always seen to proceed in the direction away from the centromere, and proximity bias effects terminate at centromeres. It is well-established in medical genetics that the short arms of acrocentric chromosomes 13, 14, 15, 21, and 22 are nonessential; indeed, in individuals bearing unbalanced Robertsonian translocations, the resulting paired short arms are often lost after a few cell divisions with no apparent clinical effect [Spinner2013]. Thus, the broader coverage of our analysis indicates that the earlier observation of whole-chromosome loss is likely a specific artifact of editing pericentromeric loci on acrocentric chromosomes, with truncation towards the telomere as the more typical effect.

Secondly, we here report a novel form of confounding for CRISPR-Cas9 phenotypic screens, which have become a powerful tool for deconvolution of biological pathways by genome-scale functional genetics [Shalem2015], and suggest a control and correction strategy. We name this effect “proximity bias”, in which the phenotypes of genes targeted using Cas9 are systematically more similar to other genes on the same chromosome arm as opposed to those on the other arm of the same chromosome or different chromosomes altogether. This confounding replicates across cell types, CRISPR delivery mechanisms, and even readout technology, with proximity bias appearing in both imaging-based phenomics assays and the viability-based screen performed in DepMap. While we did not yet have access to a genome-scale CRISPR-Cas9 experiment measured by single-cell RNA sequencing, we did observe probable truncations in smaller-scale sequencing datasets, suggesting that these screens may also be confounded. Fortunately, a simple correction method based on subtracting the mean representation of the corresponding chromosome arm from the phenotype vector of each gene is able to largely mitigate proximity bias while preserving known biological relationships. This correction depends on the presence of guides targeting non-expressed genes in the experiment as a means to estimate the undesired effect, and therefore additionally suggests that such controls are important to include in large-scale phenotypic screens. However, this method requires genome-scale datasets to derive accurate chromosome arm representations to mitigate proximity bias; thus, this bias may be difficult to correct for in smaller-scale work.

A third critical conclusion from our work is that undesired on-target effects from CRISPR-Cas9 editing are highly general across cell types, particularly including primary human somatic cells, raising potential concerns for therapeutic gene editing. Prior literature has variously examined cell lines [Cullot2019, Leibowitz2021, Przewrocka2020] or zygotes, embryos, or embryonic stem cells [Adikusuma2018, Kosicki2018, Papathanasiou2021, Alanis-Lobato2021, Zuccaro2020]. However, as the DNA damage response can vary across cell types (most obviously in cancer cells which may lack particular tumor suppressors), establishing the significance of these effects in somatic primary cells is important. Earlier work [Cullot2019] identified chromosomal truncations in cell lines and in immortalized *TP53-*null primary cells, but not in primary cells with functional *TP53*; that work, however, was limited by the limit of detection of the FISH assay used. More recent work [Tsuchida2023] has shown recurrent partial or whole chromosome-loss in *ex vivo* edited human T cells by more sensitive methods (droplet digital PCR and single-cell sequencing), and suggested that editing protocols inducing *TP53* expression prior to editing may be protective for loss. Here, we demonstrate the sensitivity of phenomics to identify that primary endothelial cells are also subject to chromosome truncation across target sites in the genome. Whereas under *ex vivo* editing conditions *TP53* induction may be feasible, our results suggest the possibility for *in vivo* editing to subject either target or bystander cells to undesired on-target effects. In particular, somatic loss of *TP53* has been observed as a recurrent phenomenon increasing in frequency with age in a variety of non-malignant tissues, including colonic epithelium and blood [Lee-Six2019, Jaiswal2019, Kessler2021], suggesting that potential risks related to *in vivo* CRISPR-Cas9 editing may be age-dependent. Further research is necessary to detect and quantify the presence of chromosomal losses in *in vivo* editing to maximize patient safety.

Based on transcriptomic evidence of truncation and genetic evidence from proximity-bias-focused analysis of DepMap, we have hypothesized a mechanism by which CRISPR-Cas9-induced truncations are introduced into a cell population. In the proposed mechanism, mitosis is an essential component for proximity bias, as the loss of large segments of chromosome is mediated by failure to propagate in daughter cells, rather than by intracellular exonuclease activity. The proposed mechanism induces testable hypotheses that may be subjects for future research, including testing the effects of mitogens, cell cycle inhibitors, or repeated passaging of edited cell populations on apparent deletion rates or proximity bias. It further suggests that functional genomics screens making use of CRISPR-Cas9 in highly mitotic cell types may be more confounded by proximity bias than those in slowly- or non-dividing cells.

From a technology perspective, our results suggest new directions for applications of Cas9-editing and improved Cas9-related protocols. Examination of proximity bias in genome-wide CRISPR maps constitutes a powerful, fine-grained tool to discover non-reference genome structure, as shown by the recapitulation of the chr5:chr19 interchromosomal fusion in U2OS. While molecular methods for translocation detection and genome assembly continue to improve, long-read sequencing and library preparation remain costly and laborious. Examination of patterns of CRISPR proximity bias may offer a simpler method to discover high-level genome architecture, particularly in genomically aberrant or unstable samples such as those found in cancers.

Finally, beyond the geometric correction developed in this work, a wide range of mitigation strategies may be developed to combat this effect. From a biological or biochemical perspective, it is likely that the use of non-cutting perturbations, e.g., CRISPRi, CRISPRoff, base editors, or RNA-targeting perturbations like Cas13d circumvents proximity bias. With cutting-based CRISPR assays, activation of p53 or DNA repair pathways (e.g., through nutlin pretreatment [Vassilev2004]) may mitigate this effect, as may the addition of free nucleotides, optimizing timing of experimental steps [Wienert2022] or extending the 5’ end of sgRNAs with cytosine bases [Kawamata2023]. Additionally, post-experimental execution strategies may also prove fruitful provided that the necessary controls are used (e.g. guides targeting non-functional sequence). Ideas here include filtering of subsets of cells in transcriptomics or patches of images in phenomics, introducing additional geometric corrections in the embedding space or utilizing loss functions during neural network training to ignore populations of affected cells. While each method has particular limitations (e.g., durability, specificity, computational intensity), the quantification methods presented in this study can be used to judge the effectiveness and drive innovation in this area.

## Limitations of the Study

A key limitation of our study is its limited molecular follow-up. While we identified truncations by transcriptomics, it is likely that our discovery power was highly limited by the intrinsic variability of RNA-seq and limited read depth. Deeper molecular characterization has been performed by other labs, and our unique contribution is breadth and scale of mapping this effect. Additionally, explicit genome-wide maps were only analyzed in two cell types – HUVEC and U2OS – and it is likely that the rate of chromosomal truncations (and the propensity across the genome) varies by cell type. Nevertheless, analysis of viability information in the 1,078 cell lines in DepMap demonstrates that this effect is not limited to HUVEC and U2OS, but likely broadly distributed across human cell types.

## Acknowledgments

We thank Hayley Donnella, Seyhmus Guler, Timothy Ahfeldt, Yolanda Chong, and Kirk Thomas for their help and discussions in generating this manuscript and the incredible Recursion lab team for design and execution of experiments.

## Author Contributions

IH contributed writing - initial draft; all authors contributed writing-review & editing; NL, CT, IH, SC, JI, LC, JJ, JU, MF, GR, and WB contributed formal analysis, visualization, validation, and methodology, CG and IH supervision, GR and WB contributed initial conceptualization.

## Data Availability

Raw images, partially-blinded metadata, and deep-learning-derived embeddings for rxrx3 are available at https://rxrx.ai.cpg0016 is available as part of the JUMP Cell Painting datasets available from the Cell Painting Gallery on the Registry of Open Data on AWS (https://registry.opendata.aws/cellpainting-gallery/). Single-cell RNA-seq datasets are available through scPerturb as described in the Methods. DepMap data are available at https://depmap.org/portal/download/all/.

## Methods

### Cell culture

HUVEC umbilical vein endothelial cells (Lonza, C2519A) at early passage are expanded within an acceptable window of in vitro culture in single use bioreactor systems that provide a 250,000 cm^2 of growth surface. This results in a yield of 10 × 10^9 cells to screen up to 4000, 1536-well plates. HUVEC are produced and banked in vapor phase liquid nitrogen and successfully seeded into high-throughput screens directly from stasis post editing. HUVEC are seeded into 1536-well microplates (Greiner, 789866) via Multidrop (Thermo Fisher) and incubated at 37 °C in 5% CO2 for the duration of the experiment.

### CRISPR-Cas9 editing

Custom-designed Alt-R CRISPR-Cas9 reagents were purchased from Integrated DNA Technologies, Inc. (IDT) and prepared following the manufacturer’s guidelines and protocols (Alt-R CRISPR-Cas9 crRNA, Alt-R CRISPR-Cas9 tracrRNA cat #1072534, Alt-R S.p. Cas9 Nuclease V3, cat #1081059). Alt-R CRISPR-Cas9 crRNA was duplexed to Alt-R CRISPR-Cas9 tracrRNA and then combined with Alt-R S.p. Cas9 Nuclease V3, following IDT guidelines, to form a functional CRISPR-RNP complex. This CRISPR-RNP complex was transfected into cells with a proprietary lipofection-based process for high-throughput application.

To control for and filter non-proximal off-target effects of individual guides, each gene was targeted with 4-12 non-overlapping guides (89% of genes targeted by 6 guides), for a total of 101,029 guides. Each guide was assessed independently in an arrayed format typically with 24 total replicate wells per guide across two executional batches.

### Phenomic imaging

Plates were stained using a modified cell painting protocol [Bray2016]. Cells were treated with mitotracker deep red (Thermo, M22426) for 35m, fixed in 3-5% paraformaldehyde, permeabilized with 0.25% Triton X100, and stained with Hoechst 33342 (Thermo), Alexa Fluor 568 Phalloidin (Thermo), Alexa Fluor 555 Wheat germ agglutinin (Thermo), Alexa Fluor 488 Concanavalin A (Thermo), and SYTO 14 (Thermo) for 35 minutes at room temperature and then washed and stored in HBSS+0.02% sodium azide. The images were acquired with ImageXpress Micro Confocal (IXMC) microscopes (Molecular Devices) in wide field mode using a PlanApo 10X 0.45 NA objective and Spectra-3 LED light engine (Lumencor). For the sake of acquisition speed, 6-channel imaging was accomplished using three combinations of two dichroic mirrors and three emission filters.

### Phenomic analysis

All images were uploaded to cloud storage and featurized by embedding them with a trained neural network using Google Cloud Platform. This network is based on the convolutional neural network DenseNet-161 [Huang2017]. We adapt this network in the following ways. First, we change the first convolutional layer to accept image input of size 512 × 512 × 6. Like DenseNet-161, we use Global Average Pooling to contract the final feature maps, which in our case are tensors of dimensions 16 × 16 × 2,208, to a vector of length 2,208. However, instead of following immediately with a classification layer, we add two fully-connected layers of dimension 1,024 and 128, respectively, and use the 128-dimensional layer as the embedding of the image. Additionally, we adapt the network to batch effects by replacing Batch Normalization layers with AdaBN [Li2018]. The weights of this network were learned by adding a classification layer to the embedding layer, with softmax activation which was optimized by training the network to recognize perturbations in the public RxRx1 dataset [Sypetkowski2023] and in a proprietary dataset of immune stimuli in various cell types.

### Generation of gene-level representations for rxrx3 HUVEC data

For this screen Recursion ran 176 12-plate experiments in 1536-well plates generating 24 images per guide for 101,029 guides and 17,065 genes. Embedding vectors for each image are centered on a set of perturbation controls, aligned using Typical Variance Normalization and aggregated to the gene-level as described in [Celik2022]. The externally released version of this data described in [Fay2023] contains all the same gene guides, but was processed with an older pipeline and contains fewer replicates per guide (18), so there may be small discrepancies with the data shown here.

### Generation of gene-level representations for cpg0016 U2OS data

Well level aggregated CellProfiler profiles were downloaded from the Cell Painting Gallery [Chandrasekaran2023]. The “Image” CellProfiler and “ObjectNumber” features were discarded and the remaining features were normalized by plate. Principal component analysis was performed using a 98% variance cutoff to reduce the dimensionality of the data followed by an additional plate normalization step. Experimental replicates were aggregated by taking the mean to yield a feature representation per gene.

#### Normalization of cosine distributions across maps

In order to make different heatmaps (e.g. rxrx3 data and cpg0016 U2OS data) visually comparable, cosine similarity values for each map were quantile normalized to a normal distribution with mean zero and standard deviation 0.2 for display purposes only.

#### Benchmarking of known relationships

To assess how well a map embedding recapitulates known biology, we calculated recall measures on known pairwise relationships from annotated sources (Reactome, HuMAP, and CORUM) as follows. Given pairwise cosine similarities between the aggregated perturbation embeddings of all perturbed genes, we select the top 5% and bottom 5% of gene pairs (excluding self relationships) from the cosine similarity distribution as “predicted relationships”. We then calculate the recall as the proportion of these predicted relationships over all relationships in the annotation source. If annotated relationships were spread randomly throughout the cosine distribution, this would produce a recall of 0.1, so that value is used as a baseline.

### Brunner-Munzel Proximity Bias Metric

In order to quantify the level of proximity bias for cell line splits in the DepMap data, we compute the effect size of intra arm relationships being larger than inter arm relationship, according to a bootstrap of the Brunner-Munzel test [Brunner2000]. In particular, we compute

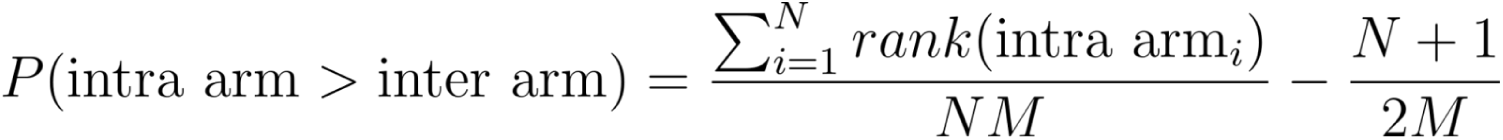

where N is the number of intra arm samples, M is the number of inter arm samples, and rank(x) is the index of sample x when all samples are sorted with ties being assigned their average rank. In order to perform bootstrapping, we set N=M and repeatedly sample N random pairs of genes from both the intra-arm and inter-arm populations for T trials. Within each of these trials we compute the cosine similarity for each gene pair and use these to compute the probability. Except where otherwise noted, we utilize N=500 and T=100. For the final score, we take the empirical mean over these trials.

### Quantification of chromosome arm bias

To quantify proximity bias at the genome-level, the Brunner-Munzel test statistic was computed between the full inter-arm and intra-arm cosine similarity distributions across all chromosome arms (not bootstrapped). The statistic estimates the probability that the intra-arm cosine similarity is greater than the inter-arm cosine similarity. For arm-level metrics we restrict the distributions to gene pairs within a given arm versus pairs with one gene on that arm.

Gene-level proximity bias quantification was computed by the Brunner-Munzel test between all cosine similarities of the gene to all other genes on the same arm and the cosine similarities to genes on other arms (not bootstrapped). The correlation between proximity bias and rank gene location between the telomere and the centromere was computed using the Spearman rank correlation of the Brunner-Munzel statistics and the ordered position of the genes on the chromosome arm.

### Analysis of public scRNAseq data

Files containing scRNAseq AnnData objects for two CRISPR-Cas9 and three CRISPRi screens were downloaded from <u>zenodo.org/record/7416068</u>: PapalexiSatija2021_eccite_arrayed_RNA.h5ad [Papalexi2021], FrangiehIzar2021_RNA.h5ad [Frangieh2021], ReplogleWeissman2022_rpe1.h5ad [Replogle2022], TianKampmann2021_CRISPRi.h5ad [Tian2021], and AdamsonWeissman2016_GSM2406681_10X010.h5ad [Adamson2016] as harmonized by the scPerturb study [Peidli2022]. Each dataset was loaded using the *scanpy* package [Wolf2018] and determined the CNV events using the *infercnvpy* package which is a scalable implementation of *inferCNV* of the Trinity CTAT project [Tickle2019]. For each of the datasets, we identified genes which, when perturbed, lead to chromosomal loss proximal to the target site (Table 1-2). A perturbed gene is called resulting in proximal chromosomal loss in a cell if (i) 70% or more of the neighboring 150 genes in the same chromosome are lost (i.e., inferred CNV value <= -0.05) in that cell. It is critical to ensure that the loss in a targeted locus results specifically from the perturbation in that locus and that it is not a promiscuous loss commonly observed when targeting other genes in distal chromosomes. Therefore, we call an observed proximal loss when targeting a gene G to be significant and specific if the fraction of cells with chromosomal loss G when perturbing G is at least three standard deviations away from the mean fraction of cells with chromosomal loss around G when perturbing any gene in the dataset. For each of the genes identified using this process, the fraction of impacted cells is reported (Table S2).

### Analysis of bulk RNAseq data

Illumina reads were aligned to the hg38 reference and gene-level counts generated for each sample using the gencode_v33 gene annotation set, and stored in AnnData objects together with sample perturbation metadata using *STAR v2.7.7a* [Dobin2013] and *scanpy* [Wolf2018]. CNV events and determination of chromosomal loss near the on-target cut site were determined using the method described above for the scRNAseq analysis. A total of 45 intron-cutting CRISPR guide perturbations were tested in HUVEC cells using this method (Table S1).

### Analysis of DepMap data

Three datasets from DepMap (https://depmap.org/portal) were analyzed: CRISPR-Cas9 22Q4 (Chronos pipeline), CRISPR-Cas9 19Q3 (CERES pipeline) and shRNA (DEMETER2 pipeline) [Dempster 2021, Meyers2017, McFarland2018]. For each gene, we treated the dependency scores across different cell lines as a feature vector and computed the cosine similarity to other genes in the dataset (Figure 3a). In order to reduce the bias towards essential genes from cosine similarity computation, we recentered the dependency scores for each gene by subtracting the mean from all cell lines. Then the cosine similarity values were quantile normalized to a Normal distribution with mean zero and standard deviation 0.2.

Additionally, we performed an analysis of the difference in proximity bias effect observed in wild type cell lines as compared to amplification or loss-of-function cell lines using the 22Q4 DepMap data. This was further stratified by looking at both a TP53 wild type background and a TP53 partial loss-of-function background. This was performed by subsetting to the cell lines that are TP53 wild type / partial loss-of-function (copy number <= 1.5) before computing the proximity bias score. In order to select the cell lines matching loss-of-function or amplification, we first subset to cell lines that have a mutation in the gene in question (where a mutation is (A) a missense, nonsense, splice-site, or stop loss mutation, (B) an inframe, start codon or frame-shift indel, or (C) a start codon SNP). We then further subset to cell lines that have copy number <= 1.5 or >= 2.5 for loss-of-function and amplification respectively. We then compute a bootstrapped version of the Brunner-Munzel proximity bias metric described above, where during bootstrapping we take a random sample of size P of cell lines, and then take the empirical mean over these 8 bootstraps. P is defined as 80% of the minimum of the number of wild type and mutant cell lines for each gene. All genes in which P < 10 are not considered.

Once this bootstrapped metric was computed for all 598 genes with a sufficient number of cell lines to meet the above conditions, we compute the difference between the wild type and each mutant condition in each background condition, and then present at the 10 genes with the highest and lowest scores for each condition (in addition to TP53 for across all cell lines), for a total of 77 unique genes across 81 conditions (Table S3).

Gene set enrichment was conducted for GO Biological Process with <u>ShinyGO 0.77</u> using the top 10 genes for the highest and lowest scores for each condition, all default setting were used (FDR cutoff 0.05, # of pathways to show, pathway size: Min 2, Max 2000, Select by FDR sort by Fold Enrichment) [Ge2020].

### Geometric method for proximity bias reduction

For each chromosome arm, first the mean vector for unexpressed genes on that arm was calculated and then that vector representation was subtracted from each gene representation on that arm. Gene locations were identified by NCBI RefSeq transcript locations against the hg38 reference assembly. Unexpressed genes were defined as those with zFPKM < -3.0 in normalized bulk RNA-seq of the given cell type [Hart2013].

## Supplemental Information

**Figure S1.**
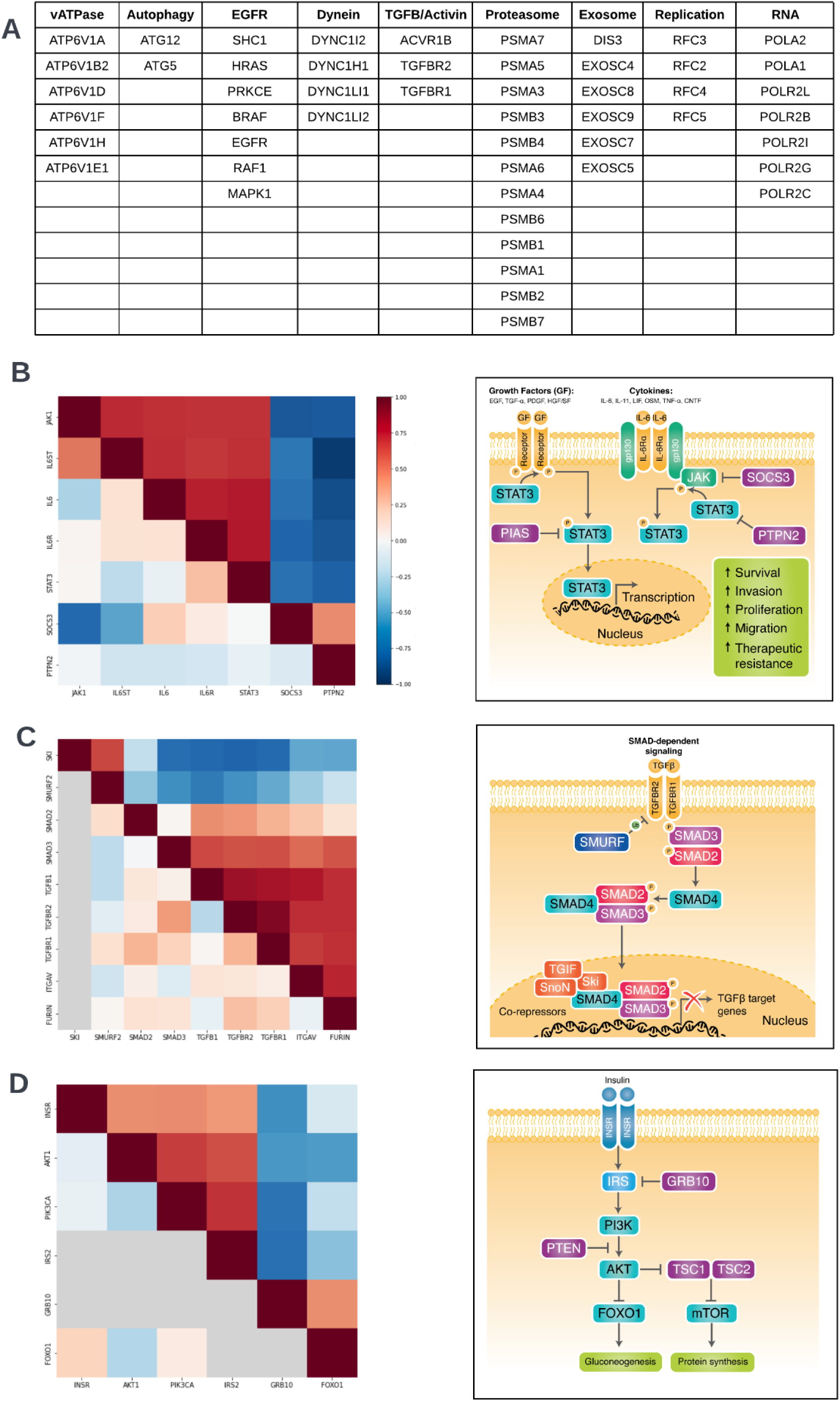
A) Table of genes shown in figure 1b B,C,D) Heatmaps of rxrx3 (above diagonal) and cpg0016 (below diagonal) data for selected biological pathways with corresponding pathway diagrams for JAK/STAT, TGFbeta and Insulin biology. Data not present in cpg0016 shown in gray. B) Example IL6 pathway: *IL6*, *IL6R*, *IL6ST*, *JAK1* and *STAT3* activate the IL-6 signaling pathway [Babon2014]. CRISPR-Cas9 targeting of these genes leads to similar cellular phenotypes and produces positive cosine similarities (red squares between *IL6*, *IL6R*, *IL6ST*, *JAK1* and *STAT3* in the heatmap). As inhibitors of the IL-6 pathway, *SOCS3* and *PTPN2* demonstrate a negative cosine similarity to the pathway components *IL6/IL6R/IL6ST/JAK1/STAT3*, especially in the rxrx3 HUVEC data (blue squares) [Babon2014, Yamamoto2002]. C) Example TGF-beta pathway: *FURIN*, *TGFB1*, *TGFBR1*, *TGFBR2*, *SMAD2*, and *SMAD3* activate the TGF-beta pathway. CRISPR-Cas9 targeting of these genes gives a similar cellular phenotype and high cosine similarity (red squares in the heatmap) [Tzavlaki2020]. *SMURF2* and *SKI* inhibit the TGF-beta pathway and CRISPR-Cas9 targeting of *SMURF2* and *SKI* show high cosine similarity to each other, but negative cosine values to *FURIN*, *TGFB1*, *TGFBR1*, *TGFBR2*, *SMAD2*, and *SMAD3* (blue squares) [Tzavlaki2020, Tecalco-Cruz2018]. Grey squares indicate genes not present in the cpg0016 data. D) Example insulin pathway: *INSR*, *IRS2*, *AKT1*, *PIK3CA* transmit insulin signaling [Haeusler2018]. CRISPR-Cas9 targeting these factors gives similar phenotypes and therefore are highly cosine similar (red squares between *INSR/IRS2/AKT1/PIK3CA* in the heatmap). *GRB10* and *FOXO1* inhibit insulin signaling, reflected in negative cosine similarities between CRISPR-Cas9 targeting of *GRB10* and *INSR*, *IRS2*, *AKT1* and *PIK3ICA* (blue squares) [Haeusler2018]. Grey squares indicate genes not present in the cpg0016 data.

**Figure S2.**
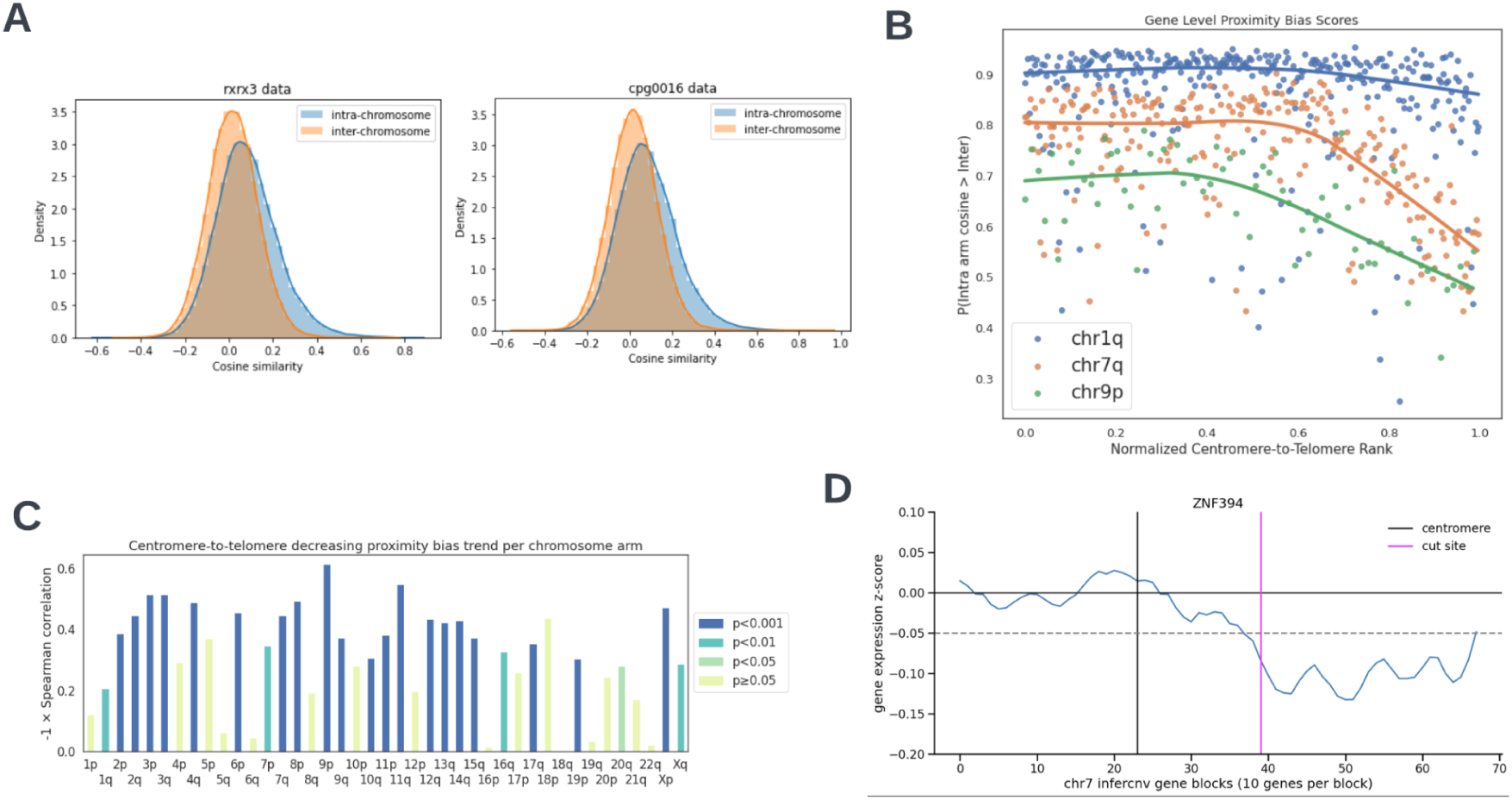
A) Distribution plots of within-chromosome and between-chromosome cosine similarity for rxrx3 and cpg0016 data. The within-chromosome distribution is shifted toward the positive which was the initial indication that some bias was present. B) Scatterplot of gene-level Brunner-Munzel probabilities versus relative chromosome-arm position for three chromosome arms in the cpg0016 dataset. The value on the y-axis estimates the probability of an intra-chromosome arm relationship involving a given gene having a higher cosine similarity than an inter-chromosome arm relationship involving the same gene. C) Spearman correlations in plots similar to C across all chromosome arms for the cpg0016 data. The height of the bar for each arm agrees well with the degree of fading in diagonal blocks in Figure 1C below the diagonal. P-values show significance of the correlation with Bonferroni correction. D) Bulk RNAseq gene count depletion for cells treated with a *ZNF394*-targeting guide relative to untreated cells in ten-gene blocks across chromosome 7. Decreased expression is evident from on the telomeric side of the cut site.

**Figure S3.**
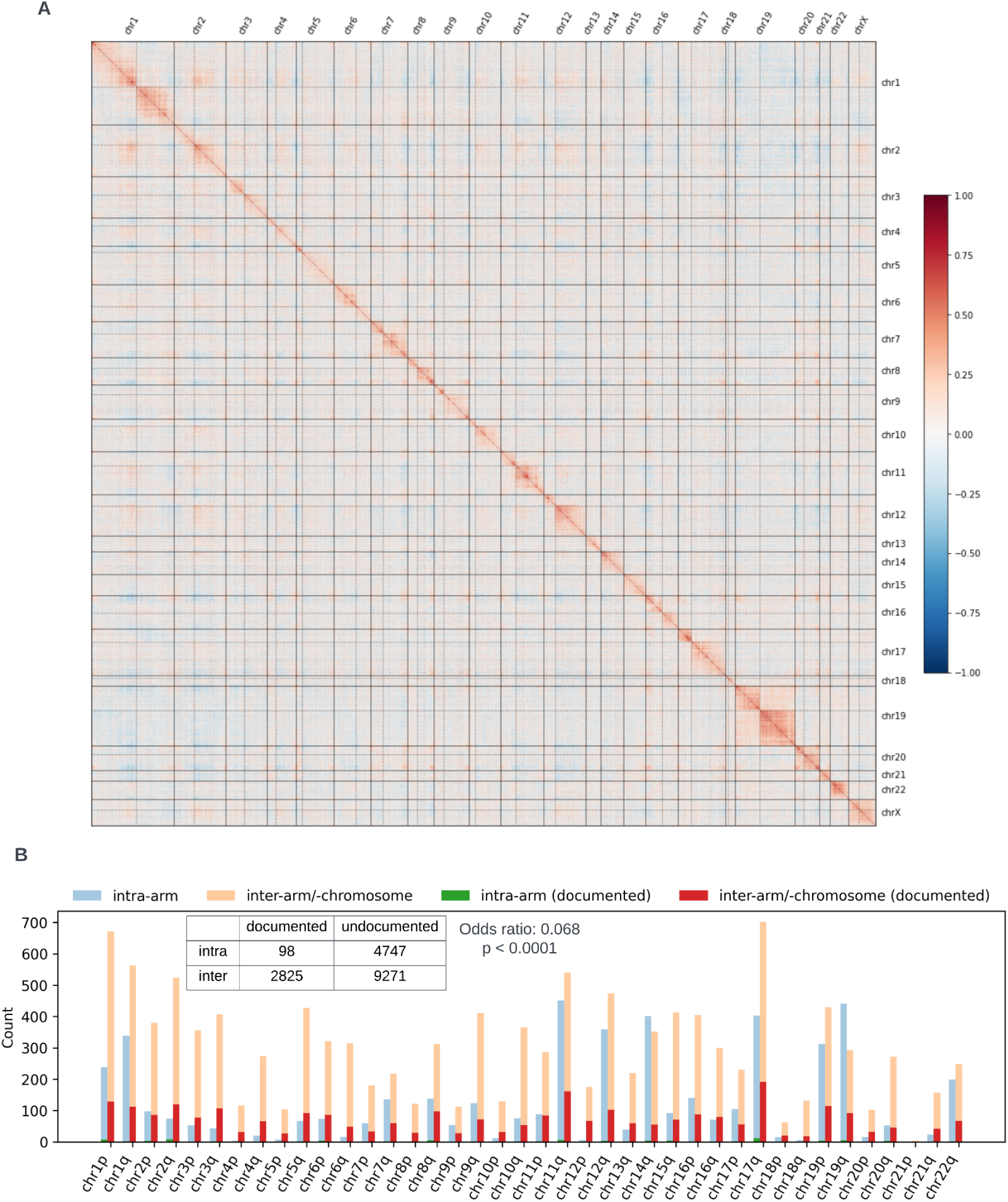
A) Genome-wide heatmap of DepMap CRISPR data processed with a different data processing pipeline, incorporating 1,078 cell lines, (DepMap 22Q4 version, Chronos pipeline) confirms the proximity bias effects [Dempster2021]. B) Counts of gene-gene relationships within and between chromosome arms for shinyDepMap 19Q3 data (blue and tan) and public annotation sets (Reactome, HuMAP and CORUM) (green and red) [Gillepsie2022, Drew2021, Giurgiu2019]. DepMap predicts a much higher proportion of within-chromosome arm relationships than are found in public annotation sets.

**Figure S4.**
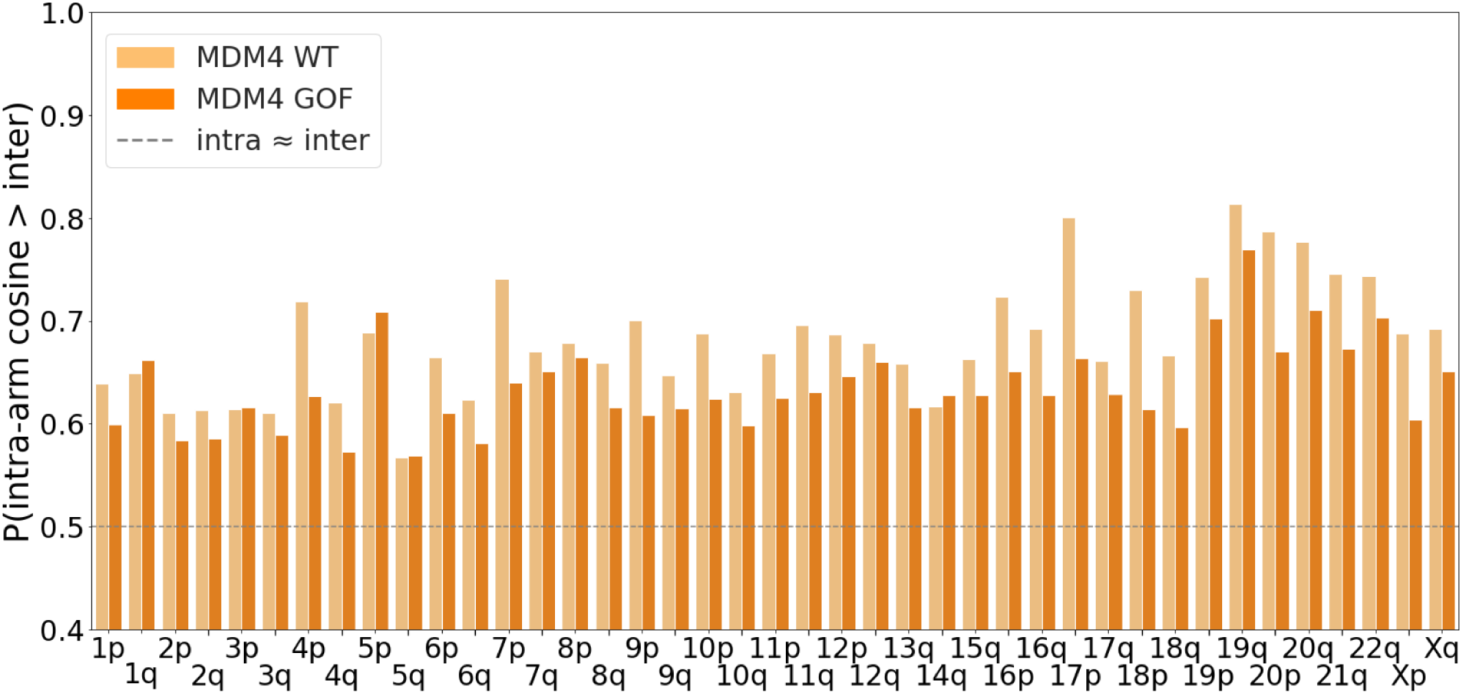
A) Barplot of chromosome arm-level proximity bias quantification by Brunner-Munzel test statistic for *MDM4* wild type (WT) and gain of function (GOF) in TP53 loss-of-function cell lines from the DepMap 22Q4 data. *MDM4* shows reduced proximity bias in most chromosome arms in cell lines with increased copy number (GOF).

**Figure S5.**
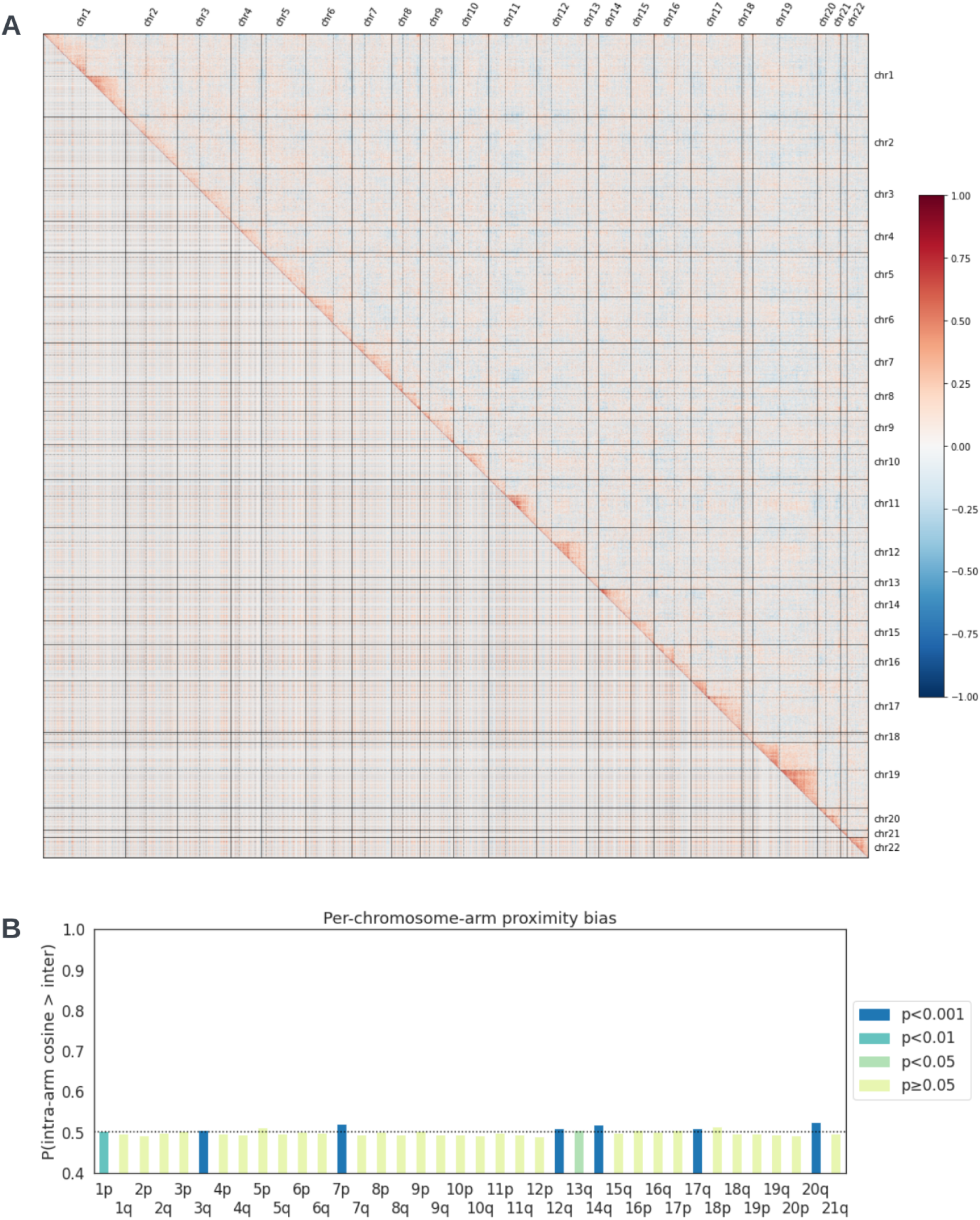
A) Split genome-wide heatmap built from 625 CRISPR cell lines, 190 shRNA cell lines and 11,169 genes shared between CRISPR and shRNA datasets in the DepMap 19Q3 data. CRISPR-Cas9 data is shown above the diagonal and shRNA data below. No proximity bias signal is visible in the shRNA data. B) Quantification of proximity bias in the DepMap shRNA dataset with Bonferroni-corrected p-values from the arm-level Brunner-Munzel test. Only a few chromosome arms display significant deviation of intra- versus inter-chromosome-arm similarities. Contrast CRISPR data from Figure 3b.

**Table S1:** CRISPR-Cas9 perturbations in HUVEC measured by bulk RNA-seq, with the fraction of wells per perturbation that exhibited specific copy loss telomeric or centromeric to the target site.

|  |  | Fraction of wells with loss<br>in given direction with<br>respect to target site |  |  |  | Fraction of wells with loss<br>in given direction with<br>respect to target site |  |
| --- | --- | --- | --- | --- | --- | --- | --- |
| gene | arm | telomeric | centromeric | gene | arm | telomeric | centromeric |
| *CEP85* | chr1p | 0.02 | 0.01 | *PCMTD1* | chr8q | 0.02 | 0.00 |
| *RCAN3* | chr1p | 0.16 | 0.04 | *HDHD3* | chr9q | 0.00 | 0.00 |
| *CD101* | chr1p | 0.00 | 0.00 | *STIM1* | chr11p | 0.00 | 0.00 |
| *RNF220* | chr1p | 0.04 | 0.00 | *KBTBD3* | chr11q | 0.02 | 0.00 |
| *EFCAB14* | chr1p | 0.00 | 0.04 | *MMAB* | chr12q | 0.00 | 0.02 |
| *SLC30A7* | chr1p | 0.07 | 0.00 | *CRY1* | chr12q | 0.00 | 0.02 |
| *GSTM4* | chr1p | 0.00 | 0.00 | *LRP10* | chr14q | 0.02 | 0.00 |
| *USP1* | chr1p | 0.00 | 0.00 | *RASGRP1* | chr15q | 0.02 | 0.00 |
| *NEXN* | chr1p | 0.02 | 0.02 | *DNAJA4* | chr15q | 0.00 | 0.00 |
| *XCL2* | chr1q | 0.00 | 0.02 | *ADAMTS7* | chr15q | 0.00 | 0.00 |
| *AVPR1B* | chr1q | 0.00 | 0.00 | *DNAAF4* | chr15q | 0.00 | 0.00 |
| *MGAT5* | chr2q | 0.02 | 0.00 | *ACSM1* | chr16p | 0.04 | 0.04 |
| *MYLK* | chr3q | 0.00 | 0.00 | *DNASE1L2* | chr16p | 0.02 | 0.00 |
| *F11* | chr4q | 0.00 | 0.04 | *GPT2* | chr16q | 0.00 | 0.09 |
| *PRMT9* | chr4q | 0.02 | 0.02 | *NOTUM* | chr17q | 0.00 | 0.07 |
| *PPARGC1B* | chr5q | 0.00 | 0.00 | *DDX52* | chr17q | 0.05 | 0.00 |
| *JARID2* | chr6p | 0.00 | 0.00 | *AKAP1* | chr17q | 0.00 | 0.04 |
| *DEFB112* | chr6p | 0.07 | 0.04 | *GZMM* | chr19p | 0.00 | 0.36 |
| *CCNC* | chr6q | 0.02 | 0.00 | *LGALS13* | chr19q | 0.09 | 0.00 |
| *HECA* | chr6q | 0.00 | 0.02 | *TGFB1* | chr19q | 0.21 | 0.13 |
| *ZNF394* | chr7q | 0.31 | 0.00 | *SRC* | chr20q | 0.05 | 0.04 |
| *CNPY1* | chr7q | 0.00 | 0.02 | *HCK* | chr20q | 0.09 | 0.00 |
| *CYP11B1* | chr8q | 0.00 | 0.00 |  |  |  |  |

**Table S2** (Lazar_et_al_2023_table_S2.xlsx)

Genes resulting in specific chromosome loss around the cut region for each CRISPR-Cas9 or CRISPRi dataset and direction tested (3’ or 5’).

**Table S3** (Lazar_et_al_2023_table_S3.xlsx)

Top 10 genes and GO Biological Process enrichment for the highest positive and highest negative proximity bias effect size when comparing mutant to wild type cell lines, stratified against *TP53* wild type or mutant background.

